# 3D Visualization of the Podocyte Actin Network using Integrated Membrane Extraction, Electron Microscopy, and Deep Learning

**DOI:** 10.1101/2021.01.29.428712

**Authors:** Chengqing Qu, Robyn Roth, Charles Loitman, Dina Hammad, Guy M. Genin, Jeffrey H. Miner, Hani Y. Suleiman

## Abstract

Although actin stress fibers are abundant in cultured cells, little is known about these structures *in vivo.* In podocytes of the kidney glomerulus, much evidence suggests that mechanobiological mechanisms underlie injury, with changes to actin stress fiber structures potentially responsible for pathological changes to cell morphology. However, this hypothesis is difficult to rigorously test *in vivo* due to challenges with visualization. We therefore developed the first visualization technique capable of resolving the three-dimensional (3D) podocyte actin network with unprecedented detail in healthy and injured podocytes, and applied this technique to reveal the changes in the actin network that occur upon podocyte injury. Using isolated glomeruli from healthy mice as well as from three different mouse injury models (*Cd2ap^-/-^, Lamb2^-/-^* and the *Col4a3^-/-^* model of Alport syndrome), we applied our novel imaging technique that integrates membrane-extraction, focused ion bean scanning electron microscopy (FIB-SEM), and deep learning image segmentation. In healthy glomeruli, we observed actin cables that link the interdigitating podocyte foot processes to newly described actin structures located at the periphery of the cell body. The actin cables within the foot processes formed a continuous, mesh-like, electron dense sheet that incorporated the slit diaphragms required for kidney filtration. After injury, the actin network was markedly different, having lost its organization and presenting instead as a disorganized assemblage of actin condensates juxtaposed to the glomerular basement membrane. The new visualization method enabled us, for the first time, to observe the detailed 3D organization of actin networks in both healthy and injured podocytes. Shared features of actin condensations across all three injury models further suggested common mechanobiological pathways that govern changes to podocyte morphology after injury.

## Introduction

End stage kidney disease (ESKD) is a major public health burden that in many cases arises when the specialized epithelial cells (podocytes) responsible for blood filtration in the kidney lose their function [1]. Although this loss of function is known to be associated with shape changes and eventual effacement of podocyte foot processes, the mechanobiological factors that underlie podocyte shape regulation and adhesion have not been fully characterized [2]. A challenge is that these cells should be studied in the natural environment of the kidney’s glomerular filtration barrier (GFB), which comprises an intricate epithelial layer of podocytes lying atop the thick glomerular basement membrane (GBM) and the endothelial layer of the glomerular capillary on the opposite side of the GBM. In this environment, podocytes develop tentacle-like foot processes that interdigitate with those of neighboring podocytes, forming slit diaphragms by stretching stacked zipper-like arrays of 40 nm-long protein assemblies over the intervening gaps [3]. However, due to these length scales being too small for conventional microscopy, and due further to the need to image podocytes within the context of a much larger, curved physiological structure [4, 5], the three-dimensional (3D) actin structures that give rise to the shape and function of podocytes has never before been quantified.

Given the dominant role of the actin cytoskeleton in determining the shapes of other epithelial cells [6–10], understanding these actin structures is especially critical for developing an understanding of how podocyte shape is regulated in both health and injury. The actin cytoskeleton is particularly responsive to mechanical stress, with both its structure and its adhesions to the matrix varying strongly with mechanical loading [11–14]. The mechanical environment of podocytes is thus critical, with stresses arising from blood pressure, cytoskeletal contraction, and fluid flow through slit diaphragms [15]. These factors further motivate the need to study the podocyte cytoskeleton in the context of an intact glomerulus in both health and injury.

When injured, podocytes undergo foot process effacement (FPE), a massive change in shape including loss of the intricate foot processes and the replacement of these with a sheet of cell membrane covering the GBM. Although a role of the actin cytoskeleton in these processes is evident from the large list of actin-associated genes whose mutation leads to FPE, the pathways leading to FPE in injury remain unclear [16–18].

Many studies have therefore focused on overcoming the challenges of scale and geometry to characterize podocytes. Transmission electron microscopy (TEM) and field emission-scanning electron microscopy (SEM) have revealed broadening of podocyte foot processes as they efface [19–21] and changes to endothelial cells associated with injury in specific glomerular diseases [22]. Two recently-developed techniques, block-face (BF)-SEM [23, 24] and focused ion beam (FIB)-SEM [25–28], have uncovered key aspects of glomerular and podocyte ultrastructure in 3D, including ridge-like promenades [25], the irregular GBM of Alport syndrome [23], podocyte-parietal cell interconnectivity [29], and changes to podocytes during the FPE that follows injury [27].

For the podocyte actin cytoskeleton itself, imaging has been limited to effaced podocytes due to the need to expose dorsal fibers. Early TEM images revealed an “actin mat” that appears in injured, effaced rat podocytes [30], but its function and three dimensional structure are not known. Super-resolution microscopy has revealed analogous, sarcomere-like structures of alternating myosin IIA localization and synaptopodin (with alpha-actinin-4) in effaced podocytes of mouse models of nephrotic syndrome and podocyte injury (*Cd2ap^-/-^, Lamb2^-/-^,* Alport syndrome, and adriamycin-induced nephropathy), as well as in kidney biopsies from patients with minimal change disease, focal segmental glomerulosclerosis, and diabetic nephropathy [31]. However, the relationships of these structures to podocyte health and to kidney function have not been established.

From the perspective of mechanobiology, actin is a critical mediator of cell shape and function, as well as a critical mediator of cell-matrix interactions and mechanosensing [32–35]. The challenge of imaging actin structures *in vivo* has been a factor limiting the understanding of a great many cell types, and underlying uncertainty about the role of structures such as stress fibers in the physiology of mature, healthy cells [36, 37]. There is thus a pressing need for a broadly applicable technology to image the actin cytoskeleton at high resolution. We therefore developed such a technique, and applied it to test the hypothesis that podocyte injury is associated with disruption of the podocyte’s 3D actin network.

## Materials and methods

### Animal models

C57BL/6J mice (The Jackson Laboratory) were used for isolation of WT glomeruli. Three mouse models of nephrotic syndrome and foot process effacement were used: **1**. *Cd2ap^-/-^* mice: A model of primary podocyte injury with nephrotic range proteinuria and FSGS due to slit diaphragm defects (age ~16 days) [38]. **2.** *Lamb2^-/-^* mice: A model of Pierson syndrome, a GBM disease with nephrotic syndrome (age ~16 days) [39]. **3**. *Col4a3^-/-^* mice: A model for Alport syndrome, a GBM disease with thickening/splitting of the GBM and delayed proteinuria (age ~60 days) [40]. Mice were anesthetized using Ketamine/Xylazine cocktail. Glomeruli were isolated by perfusing mice with Dynabeads as previously described [41] but without collagenase digestion.

### Glomerular isolation and membrane-extraction

Isolated glomeruli were subjected to membrane-extraction as previously described [42]. All rinses utilized a super-magnet to help pull down the pellet and prevent loss of glomeruli during washes. The extraction buffers (KHMgE buffers) contained 10 μM phalloidin, 10 μM taxol, 2 mM ATP, 1 mM phenylmethylsulfonyl fluoride, 10 μM leupeptin in 0.1 M KCl, 30 mM HEPES at pH 7.2, 10 mM MgCI_2_, and 2 mM EGTA at pH 7.1. Extraction solutions contained 0.01% saponin and 0.5% Triton X-100. Before the extraction procedure, samples were preincubated with the KHMgE solution for 1.5 minutes, and then detergent extracted for 1.5 minutes in the respective extraction solution before fixing the sample for 10 minutes in 4% PFA in KHMgE buffer in the presence of 0.01% Saponin, 0.5% Triton X-100 before fixing it further in 2% PFA / 2% glutaraldehyde in PBS at RT.

### Adherence of Glomeruli to Glass

In preparation for EM, glass coverslips were cut into 3mm x 3mm squares, cleaned in chromo-sulfuric acid solution and washed extensively with dH2O. Coverslips were then incubated in 5 mg/ml LMW poly-l-lysine in 0.1M KCl for 30 minutes and then washed 3 x 5 minutes in 0.1 M KCl. Coverslips were placed onto a parafilm coated super magnet, the residual KCl solution wicked off the top of the slip and a 10 microliter drop of suspended glomeruli in KHMgE buffer (70 mM KCl, 30 mM HEPES 5mM MgCl, 3 mM EGTA pH 7.0) was deposited onto the slip and let adhere 5 minutes at RT. Following adhesion, coverslips were transferred into 2% glutaraldehyde in KHMgE buffer for 30 minutes, then either washed 3 x 10 minutes 0.15 M Na Cacodylate with 2mM CaCl_2_, pH 7.4, and processed for SEM or washed 3 x 5 minutes in dH2O and processed by QF-DEEM. A super-magnet was utilized to help secure adhered glomeruli to the coverslip surface for solution transfers hereafter.

### SEM preparation

Isolated glomeruli were subjected to membrane extraction as previously described [42]. Samples were stained with 1% osmium tetroxide in 0.15 M Na Cacodylate with 2 mM CaCl_2_, pH 7.4 for 1 h, room temperature, in the dark. After 3 x 10-minute rinses in dH20, samples were dehydrated in a graded ethanol series (50,70, 90, 100 and 100%) for 10 minutes each and then critical point dried (Leica CPD 300). Glass slips were adhered to a conductive carbon adhesive tab on aluminum stubs. Sample stubs were then sputter coated with 6-nm iridium, (Leica ACE 600) and imaged on an SEM (Zeiss Merlin equipped with a Gemini ll electron column) at an accelerating voltage of 3 keV with a beam current of 200 nA with secondary electrons detected by an Everhart-Thornley (SE2) detector.

### Quick-Freeze Deep-etch EM (QF-DEEM)

Quick-freezing was accomplished by previously reported protocol with minor modifications [43–48]. Briefly coverslips containing extracted glomeruli were rinsed in dH2O, and frozen by abrupt application of the sample against a liquid helium–cooled copper block with a Cryopress freezing machine. Frozen samples were transferred to a liquid nitrogen-cooled Balzers 400 vacuum evaporator, fractured at −100*C, then etched for20 minutes at −80°C, and rotary replicated with approximately 2-nm platinum deposited from a 20°angle above the horizontal, followed by an immediate, approximately 10-nm stabilization film of pure carbon deposited from an 85°angle. Replicas were floated onto a dish of concentrated hydrofluoric acid and transferred with a glass rod through 3 rinses of dH20, one rinse of household bleach, then 3 additional rinses of dH20,all solutions containing a loopful of Photo-flo.Replicaswere picked up on Luxel grids (LUXEL), and photographed on a JEOL 1400 microscope with attached digital camera (AMT, AMT Imaging in Woburn, MA).

### FIB-SEM sample preparation

FIB-SEM sample preparation and imaging were performed as described [49, 50]. Samples were incubated in a fixative containing 2.5% glutaraldehyde and 2% paraformaldehyde in 0.15 M cacodylate buffer containing 2mM CaCl_2_, pH 7.4 overnight at 4°C. The samples were then stained according the methods described by Deerinck et. al [49]. In brief, samples were rinsed in cacodylate buffer 3X for 10 min each, subjected to a secondary fixation for one hour in 2% osmium tetroxide/1.5% potassium ferrocyanide in cacodylate buffer for one hour, rinsed in ultrapure water 3X for 10 min each, and stained in an aqueous solution of 1% thiocarbohydrazide for one hour. After this, the samples were again stained in aqueous 2% osmium tetroxide for one hour, rinsed in ultrapure water 3X for 10 min each, and stained overnight in 1% uranyl acetate at 4°C. The samples were then washed in ultrapure water 3X for 10 min each and *en bloc* stained for 30 minutes with 20 mM lead aspartate at 60°C. After staining was complete, samples were briefly washed in ultrapure water, dehydrated in a graded acetone series (50%, 70%, 90%, 100% x2) for 10 min in each step, and infiltrated with microwave assistance (Pelco BioWave Pro, Redding, CA) into Durcupan resin. Samples were cured in an oven at 60°C for 48 hours.

### FIB-SEM imaging

Post resin curing, samples were exposed with a razor blade, and 70 nm thick sections were prepared on silicon wafer chips. These chips were then adhered to SEM pins with carbon adhesive tabs, and large areas were then imaged at high resolution in a FE-SEM (Zeiss Merlin, Oberkochen, Germany) at 5 KeV and 3 nA using the ATLAS (Fibics, Ottowa, Canada) scan engine to tile resin sample surface and identify regions of interest for further analysis. Once regions of interest were identified, resin blocks were mounted onto SEM pins with silver epoxy and sputter coated with 6 nm of iridium (Leica ACE 600, Vienna, Austria) at a 45° angle with rotation on a planetary stage to ensure the entire block was coated.

Wildtype control, *Lamb2^-/-^* and Alport syndrome sample blocks were loaded into the FIB-SEM (Zeiss Crossbeam 540, Oberkochen, Germany), and the identified regions of interest were located by secondary electron imaging at 5 KeV and 900pA. Once a region was found, the sample was prepared using the ATLAS (Fibics, Ottowa, Canada) 3D nanotomography routine [50]. In short, a platinum pad was deposited on a 20 μm x 20 μm region of interest at 30 KeV and 1.5 nA. Three vertical lines for focus and stigmation and two angled lines for z-tracking were milled into the platinum pad at 300 pA, then filled with carbon at 50 pA to fill the tracking/alignment marks followed by an additional deposition of a protective platinum pad at 1.5 nA. A rough trench 40 μm wide and 25 μm deep was then milled at 30 nA and polished at 7 nA. Once polished, face detection, focusing, and z-tracking were all performed on the fiducial marks that were milled into the platinum pad. Serial block-face imaging was performed at 2.5 KeV and 900 pA using the SE2 and ESB detector with a grid voltage of 1050 V. The block was milled at a current of 300 pA with 10 nm slices and images were acquired at a resolution of 10 nm/pixel with a dwell of 4 μs and a line average of 5 for a total z-depth of about 15 μm. The stack of acquired images was aligned using Atlas 5 (Fibics, Ottowa, Canada).

For the *Cd2ap^-/-^* sample, the block was loaded into the FIB-SEM (Helios 5 UX DualBeam, Thermo Fisher Scientific, Brno, Czech Republic), and the identified regions of interest were located by secondary electron imaging at 5 KeV and 800 pA. Platinum was deposited on a 20 μm x 20 μm region of interest at 30 KeV and 1.2 nA, and a rough trench 90 μm wide and 35 μm deep was then milled at 47 nA using Si-multipass module and polished at 1.2 nA. Trenches on the left and right side of the platinum pad were milled at 20 nA using Si-ccs module. A SEM fiducial marker was placed on the face of a right trench parallel to the imaging face by depositing a platinum pad at 750 pA and then milling a distinctive pattern at 41 pA. A FIB fiducial marker was placed on the top side of the imaging volume by depositing a platinum pad at 1.2 nA and milling an “X” pattern at 41 pA. Serial block-face imaging was performed at 1.8 KeV and 800 pA using the ICD and TLD detector in backscattered mode and the ASV 4 software (Thermo Fisher Scientific, Hillsboro, Oregon, USA). The block was milled at a current of 750 pA with 10 nm slices and images were acquired at a resolution of 10 nm/pixel with a dwell of 5 μs and a line average of 2 for a total z-depth of about 15 μm. Stack of acquired images was aligned using Amira 2019.4 (Thermo Fisher Scientific).

### Post-imaging analysis and manual segmentation

Image processing and data analysis were performed using a Data Workstation with Dual Intel Xeon E5-2650 2.3Ghz 10-Core Hyper-threaded Processors and 256 GB DDR4-2133MHz RAM in the Washington University Center for Cellular Imaging. For manual segmentation, selected ROI were cropped from the stack using Fiji/ImageJ and segmented using Amira software (v9.3). Using an interactive tablet system (INTUOS Pen Tablet medium), the GBM, nuclei, foot processes, and podocyte cell body were selected manually on each slice then assembled into a 3D object. For both WT control and Alport syndrome datasets, 150 slices were segmented.

### Deep learning model and supervised segmentation

To segment the GBM and podocyte actin cytoskeleton from FIB-SEM image stacks of glomeruli of WT control, *Cd2ap^-/-^, Lamb2^-/-^* and Alport syndrome mice, Fiji/ImageJ (version 1.53C) was utilized with the Waikato Environment for Knowledge Analysis (Weka) trainable segmentation plugin (version 3.2.34) [51, 52]. Before training the model, features were selected. For edge detection features, the *Sobel filter, Hessian* and *Difference of Gaussian* options were selected. For texture description features, the *Structure* and *Neighbors* options were selected. For noise removal and edges detection features, *Gaussian blur* and *Membrane projections* were utilized [52].

To train deep learning models, we generated independent trained classifiers for each dataset: WT control, *Cd2ap^-/-^m Lamb2^-/-^* and Alport syndrome. For each classifier, five elements were classified and assigned to the selected region on the image stack via the graphic user panel. These five elements represented “actin”, “GBM”, “background”, “microtubules” and “gaps”, where “gaps” indicated the peripheral fading boundary between the podocyte actin cytoskeleton and the “background” in the image. The development and training of deep learning models for each of the four conditions was based on a region of interest (ROI) within FIB-SEM image stacks. The training ROI was 251 slices at 587 x 218 pixels for WT control, 251 slices at 252 x 676 pixels for *Cd2ap^-/-^*, 201 slices at 378 x 484 pixels for *Lamb2^-/-^,* and 150 slices at 286 x 172 pixels for Alport syndrome. The classifiers were trained based on the ROI and the selected features, one for each of the image stacks, except for Alport syndrome. The classifiers were then applied to the ROI to generating probability maps.

Because results revealed that the central actin cable was connected to the slit diaphragm, additional training was required to segment one from the other. For that purpose, we added one more “connection” class for the slit diaphragm structure, followed by machine training based on that plus the other five classes listed above using the procedures described above.

For Alport syndrome, because of the complexity arising from GBM splitting/thickening and actin condensations protruding into the GBM, direct training of the classifier was not possible. Thus, manual segmentation was applied first to separate the GBM and the actin condensations into two different stacks. Then two classifiers were trained for GBM and actin condensations separately. Based on the two stacks, we generated two probability maps, one for GBM and one for actin condensations. Finally, the two probability maps were combined into one final probability map.

All probability maps were projected into 3D models using Amira software (Thermo Fisher Scientific, Version 2019.4), with each probability map having one channel for each class. The final channels used for 3D visualization were the “GBM” and “actin” channels, resulting in low 3D visualizations of the structures of the GBM and podocyte actin network.

## Results

### Membrane-extraction technique and Scanning electron microscopy (SEM) imaging of podocytes

To directly observe the cytoskeletal network of podocytes, we adapted the method of cytoskeleton stabilization and membrane extraction [42]. During the membrane-extraction process (Triton and saponin at RT), we used microfilament-stabilizing agents to prevent depolymerization of the actin network (phalloidin, 10 μM) and microtubules (Taxol, 10 μM). To determine the efficiency of this approach for extracting cell membranes while maintaining intact microfilament networks, we used SEM to compare intact (Fig.1a) and membrane-extracted healthy glomeruli (Fig. 1c and e). We observed a clear lack of membranes in the extracted samples, but the cytoskeletal networks within the major processes and the foot processes appeared well preserved (Fig. 1c and e, arrows, Fig. 1g and h), as were the slit diaphragms and the GBM underneath.

**Figure. 1.**
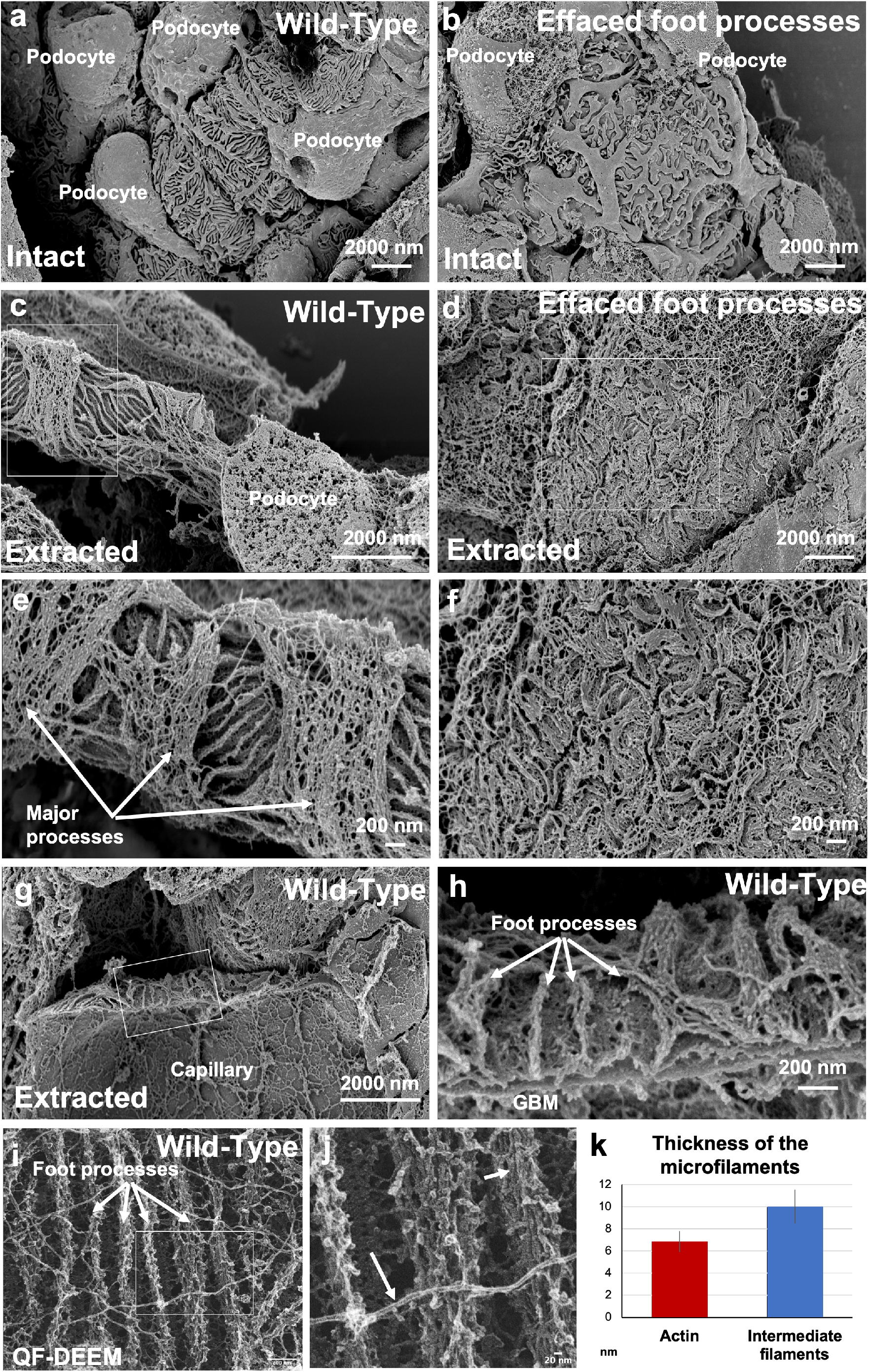
Scanning electron microscopy (SEM) of intact and membrane-extracted podocytes show the exposed actin network of the interdigitating foot processes after cell membrane removal. (**a**) SEM image of an intact healthy glomerulus shows 4 podocytes with major processes and interdigitating foot processes. (**b**) SEM image of an intact glomerulus from a *Cd2ap^-/-^* mouse with signs of foot process effacement. (**c**) SEM image of a membrane-extracted healthy glomerulus shows a capillary loop with the cytoskeletal filaments of major processes and interdigitating foot processes. Notice the podocyte nucleus in the bottom right corner. (**d**) SEM image of a membrane-extracted glomerulus from a *Cd2ap^-/-^* mouse shows an effaced area with irregularly distributed wavy actin cables covering the GBM. (**e**) Zoomed-in region from (**c**) shows the major processes (arrows) and the interdigitating foot processes (arrowheads) leaving the slit diaphragm areas in between. Note the porous nature of the slit diaphragms. (**f**) Zoomed-in region from (**d**) shows the irregularly distributed thick actin cables in the effaced areas of *Cd2ap^-/-^* podocytes. (**g**) and (**h**) SEM images of a membrane-extracted and fractured healthy capillary show the endothelial cell cytoskeleton, the GBM and foot processes (arrows) with the slit diaphragms in between (asterisks). (**i**) Quick-Freeze Deep etch EM (QF-DEEM) micrograph of a membrane-extracted healthy podocyte shows the actin bundles of foot processes (arrows in **i**) and slit diaphragms in between. Note the long microfilaments running above the foot processes. (**j**) High magnification image of the boxed area in (**i**) shows details of the actin bundles (short arrow) and the microfilaments (long arrow). (**k**) Measurements of the thicknesses of the individual filaments in the foot processes (similar to the short arrow in (**j**)) and the long filaments nearby (long arrow in (**j**)) are consistent with the first being actin and the latter being intermediate filaments.

To identify changes after injury, we used the same technique to membrane-extract glomeruli from mouse podocyte injury models (*Cd2ap^-/-^* mice, Fig. 1b, d and f; *Lamb2^-/-^* mice, Fig. S1). Intact samples showed widened foot processes as a sign of FPE when compared with the WT (Fig. 1b). Interestingly, extracted glomeruli from injury models showed thick, disorganized microfilament assemblies covering the capillary loops (Fig. 1d and f) that were not present in the WT control (Fig. 1c and d). These data suggest that the membrane-extraction process successfully dissolved podocyte cell membranes when exposed to the extraction buffer while maintaining cytoskeletal networks.

In order to gain more insights about the structures that we observed, we imaged WT membrane-extracted glomeruli using two types of electron microscopy approaches, SEM (Fig. 1g and h) and QF-DEEM (Fig. 1i and j). Comparing the two techniques showed similar results, with high levels of preservation of the cytoskeletal structures in foot processes and across the GBM. QF-DEEM platinum replicas confirmed the presence of ~7 nm-diameter filaments arranged as bundles in the foot processes, consisting of uniform globular subunits with polar orientation, which are characteristic features of assembled actin [44, 53]. Thicker filaments with a diameter of ~10 nm identified within the cytoskeleton are presumed to be intermediate filaments, based on their size, structural stability, and frequent proximity to the cell nucleus. With consideration of variable deposition of the 2 nm platinum film evaporated on our samples, resulting from the angle of evaporation and the 3D topography of the specimens, variable increases in filament diameter are expected. This variability is consistent with our diameter measurements of ~7 nm for actin and ~10 for intermediate filaments. From these observations, we conclude that the observed filaments are actin and intermediate filaments, respectively (Fig, 1k) [54].

### Focused ion beam scanning electron microscopy (FIB-SEM) imaging of membrane-extracted healthy podocytes

Because SEM only allows visualization of surface topology, we sought a method to view the whole depth of the GFB. To achieve this, we used FIB-SEM, which generates an image stack in the Z-direction, thus providing a view of the structures located deep in the tissue [27]. Imaging the membrane-extracted glomeruli with FIB-SEM showed excellent preservation of glomerular structures, including basement membranes (GBM and Bowman’s capsule) and cell nuclei (Fig. 2a, b and c; Mov 1).

**Figure. 2.**
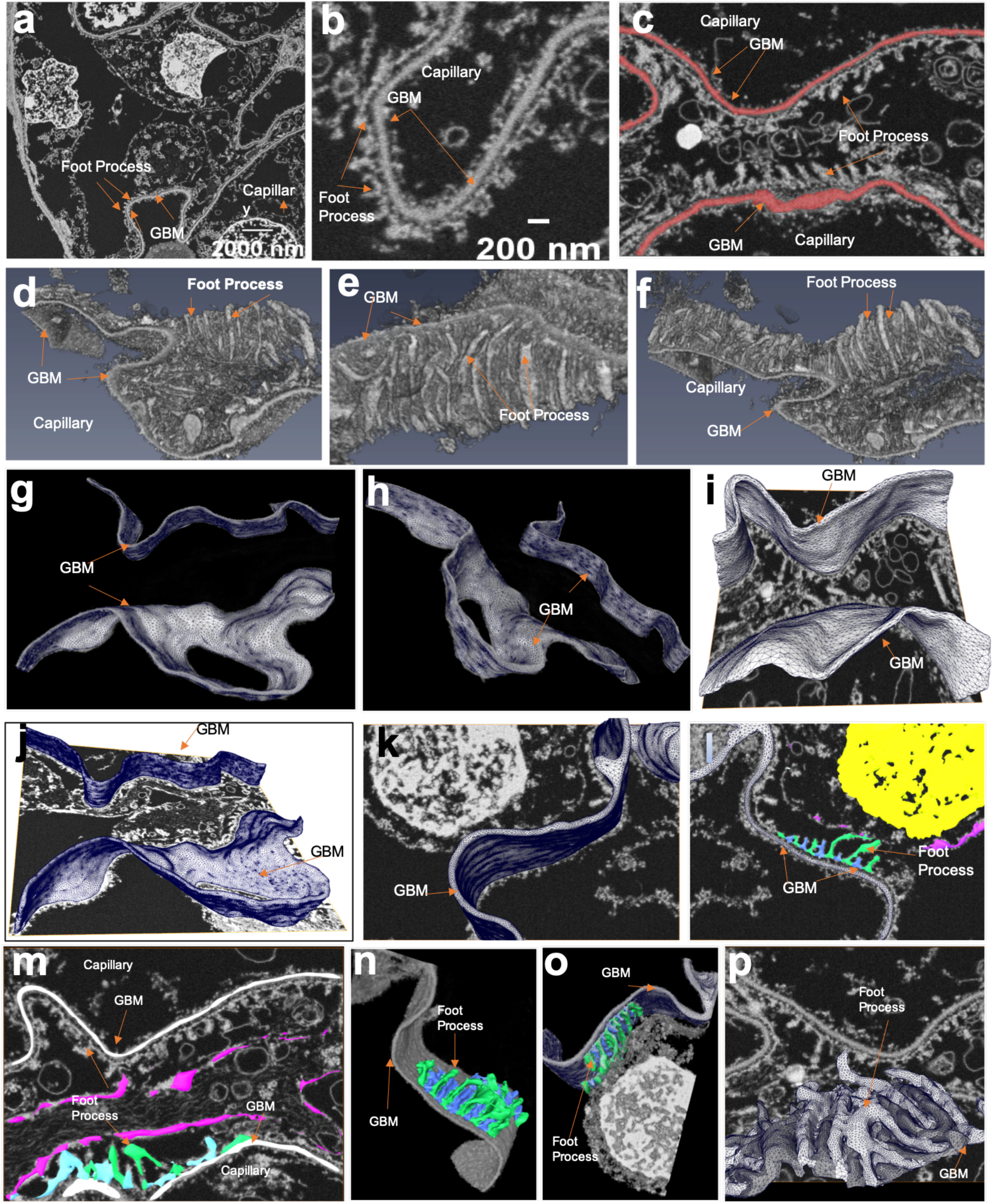
Manually segmented FIB-SEM 3D visualization of membrane-extracted healthy podocytes reveals interdigitation of foot processes. (what about: reveals the detailed architecture of the capillary wall) (**a**) FIB-SEM overview image shows a healthy glomerulus with intact nuclei and GBM. Notice the lack of cell membranes and the electron dense actin cables in podocyte foot processes. (**b**) Zoomed-in area in (**a**) shows a capillary loop with intact GBM and foot processes. Note the continuity between the foot processes and the adjacent slit diaphragms (arrows). (**c**) Region of interest (ROI) for segmentation of the actin network. The two red areas indicate the two GBMs, one in the upper area of this panel and one in the lower. (**d**) 3D top view of the capillary stack, using the GBM and foot processes in the upper ROI in panel (**c**), shows the complete GBM. (**e**) 3D front view of the same structure in panel (**d**) shows the foot processes. (**f**) 3D bottom view of the same structure in panel (**d**) shows both the GBM and the foot processes. (**g**) Top view of a 3D visualization of the GBM (red areas in panel (**c**)) by manual segmentation. (**h**) Side view of a 3D visualization of the GBM in panel (**c**) by manual segmentation. (**i**) Top view of the manually segmented 3D visualization of the GBM combined with an orthogonal image from the FIB-SEM image stack shown in (**c**). (**j**) and (**k**) Side views of the manually segmented 3D visualization of the GBM combined with panel (**c**). (**l**) and (**m**) Low and high magnification views of the segmented the actin cables of two neighboring foot processes atop of the GBM highlighted in green and blue. The nucleus is segmented as yellow and the actin patches at the periphery of the cell body are labeled in pink. (**n**) and (**o**) 3D visualization of manually segmented GBM, foot processes, and cell body. (**p**) 3D visualization of manually segmented foot processes with an orthogonal image from the FIB-SEM image stack.

Results revealed that the central actin cables in the foot processes were connected with the neighboring slit diaphragms located just above the GBM, forming a continuous structure (Fig. 1b). This was further evident from detailed analysis and 3D visualization of these podocyte structures, which showed the actin cables in the foot processes (Fig. 2b, c d, e and f). 3D visualization (Fig. 1d, e and f) as well as manual segmentation of the various structures of the GFB including the GBM (Fig.1g, h, I, j and k) and the foot processes (Fig. 1l, m, n, o and p) showed excellent preservation of the podocyte foot processes and the slit diaphragms. Collectively, this 3D visualization proves that the membrane-extraction approach faithfully preserves the structures observed by TEM and SEM and that actin cables in the foot processes and the nephrin-based slit diaphragms form one continuous structure.

### Actin cables within the podocyte cell body

One interesting observation in the membrane-extracted glomeruli was the presence of peripheral, electron-dense patches around the podocyte cell bodies (Fig. 3a and b, arrows) and the major processes (Fig. 3g, arrows). These are reminiscent of the synaptopodin-positive patches that we observed previously using STORM super-resolution imaging [31]. Surprisingly, when we segmented those patches in the FIB-SEM stacks and rendered them in 3D, they formed thick cables, running in parallel to one another along the longitudinal axes of the cell body (Fig. 3c, d, e and f) as well as around the major processes (Fig. 3g, h, i, j and k, Mov 2 and 3).

**Figure. 3.**
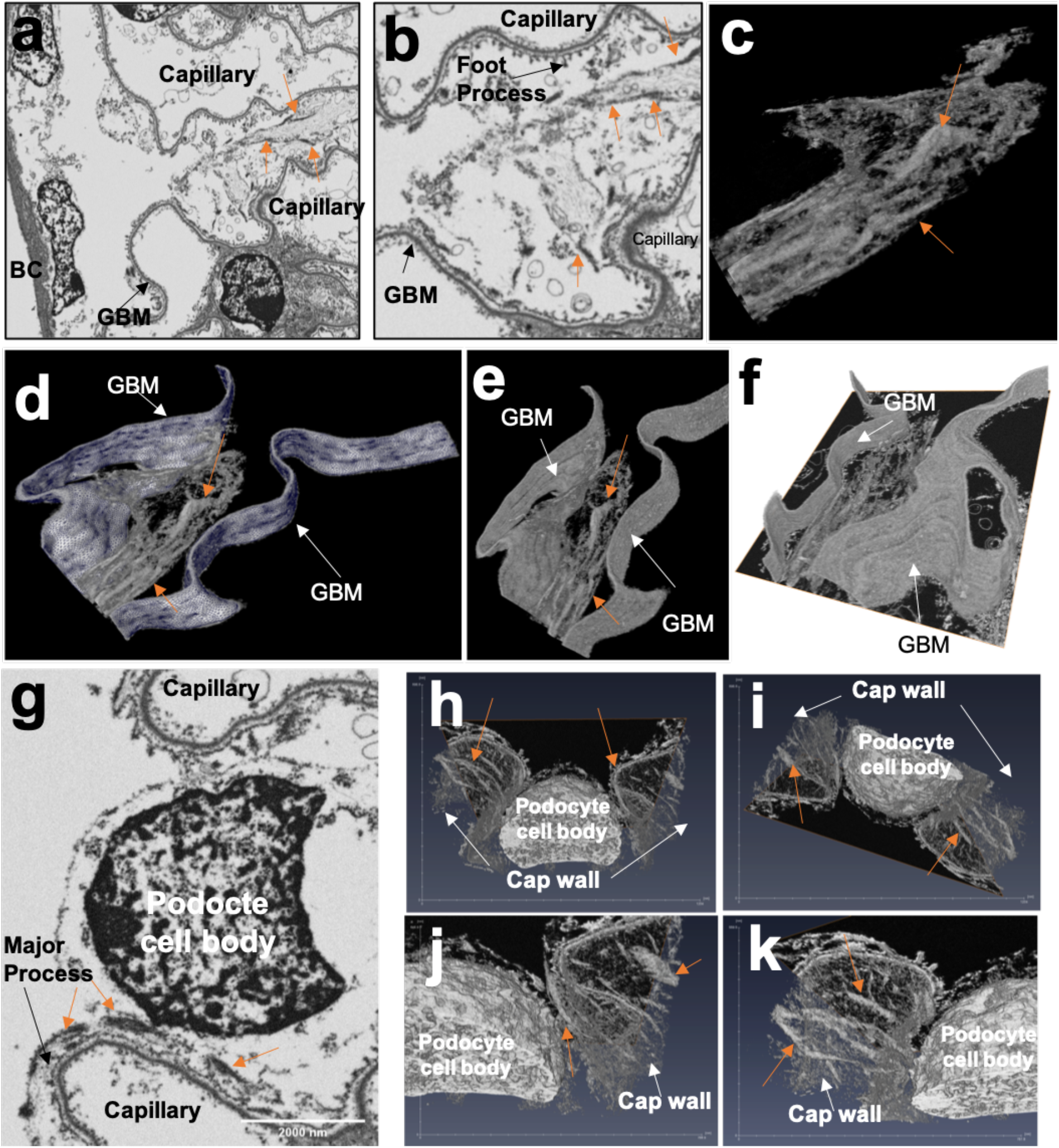
Manually segmented FIB-SEM images of a podocyte cell body of in a membrane-extracted healthy glomerulus shows peripheral thick actin cables along the cell body axis. **(a-b)** Overview **(a)** and zoomed-in (**b)** FIB-SEM single-frame image of a membrane-extracted glomerulus shows actin cables in patches (arrows) reinforcing podocyte cell bodies. **(c)** 3D segmentation of the electron dense patches around the podocyte cell body shows that they are parts of thick parallel actin cables running along the longitudinal axis of the cell. **(d**, **e** and **f)** 3D renderings of both the podocytes’ peripheral actin cables and the GBM show the position of these longitudinal cables relative to the GBM. (**g**) Overview of the segmented actin assembly around the podocyte major processes. This image is one slice of the of the segmented stack showing the actin patches around the major processes (orange arrows). Overview 3D visualization (**h** and **i**) and zoomed-in views (**j** and **k**) of the segmented areas in (orange arrows in **e**) show thick branched actin cables around the major processes (orange arrows). For orientation, white arrows indicate the opening of the capillary loops. BC: Bowman’s capsule, MP: Major processes. Scale: tickmarks in the x and y axes are in nm.

### Electron microscopic imaging of membrane-extracted injured glomeruli

To identify how the actin cytoskeleton changes after podocyte injury, we imaged the three mouse models of glomerulopathy (i.e., *Cd2ap^-/-^, Lamb2^-/-^* and Alport mice) after membrane extraction using both TEM and FIB-SEM. When compared with the WT control, TEM images of nephrotic models showed effaced areas above the GBM (Fig. 4a, b, c and d). All three models showed electron dense cytoskeletal structures juxtaposed to the GBM (red arrows), suggesting that these are part of the “actin mat” described previously [30].

**Figure. 4.**
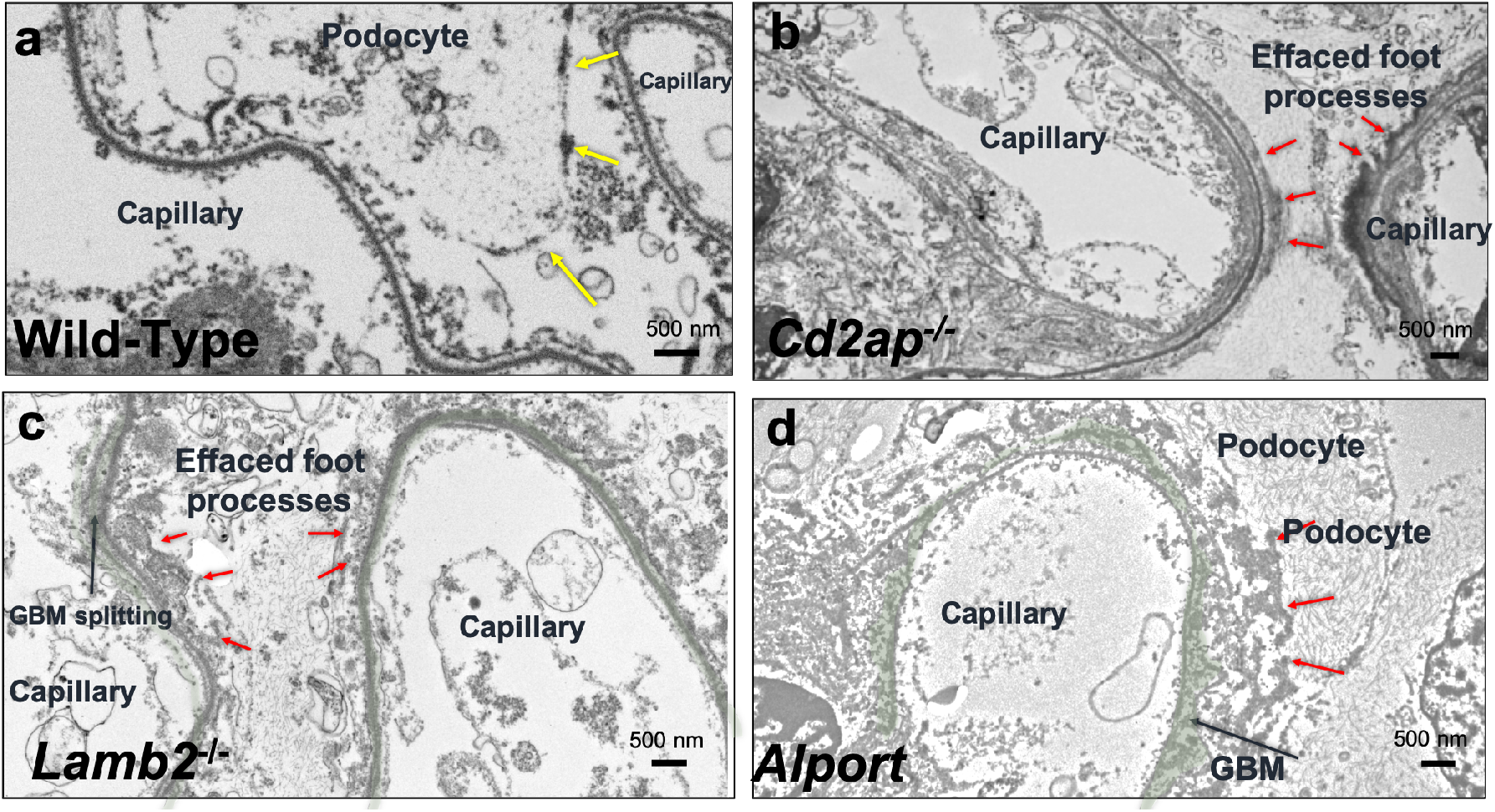
Transmission electron microscopy (TEM) images of membrane-extracted podocytes of healthy and injured glomeruli show actin condensations in the bottom of the effaced areas. **(a)** TEM image of membrane-extracted healthy podocytes show organized actin cables in the foot processes. Notice the continuity between these actin cables and the adjacent slit diaphragms. Yellow arrows indicate the peripheral actin patches around the cell body. **(b)** TEM image of membrane-extracted podocytes from a *Cd2ap^-/-^* glomerulus shows effaced foot processes and electron-dense condensations juxtaposed to the GBM (red arrows). **(c)** TEM image of membrane-extracted podocytes from a *Lamb2^-/-^* glomerulus shows effaced foot processes and electron-dense condensations juxtaposed to the GBM (red arrows). Note the GBM splitting (black arrow). **(d)** TEM image of membrane-extracted podocytes in an Alport glomerulus shows effaced foot processes and electron-dense condensations juxtaposed to the GBM (red arrows). Note the irregularly split, heterogeneous GBM (Black arrow).

### Applying a deep learning model for 3D image segmentation of healthy podocytes

To eliminate user bias in the segmentation of FIB-SEM image stacks, we applied a machine learning tool that allowed us to segment the GBM and the actin structures surrounding it using the trainable Weka plug-in in Fiji/ImageJ software [52]. In healthy glomeruli, machine learning and segmentation of the GBM in the training stack (15 frames) was straightforward, but accurate identification of the actin cables in the foot processes required multiple rounds of training (see the Methods section for details). Applying this training approach to a training stack from a set of healthy images (Fig. 5a) generated a trained model for segmentation of the GBM and the actin assembly in the foot processes (Fig. 5b) that then was applied to a larger image stack. Applying this to a FIB-SEM stack of 251 images (Mov 4) generated an extensive probability map of the two classes of interest (Fig. 5c; GBM (Red), actin in the foot processes (light green)) that subsequently was used to generate a 3D model of the GBM and the actin cables in the foot processes (Fig. 5d, e and f; Mov 5). Moreover, by adding one more class for the slit diaphragm, we trained imaging stacks from the healthy glomerulus (Fig. 5g), which allowed us to segment actin in the foot process from the slit diaphragm (Fig. 5h). The probability map generated (Fig. 5i; GBM (Red), actin in the foot process (light green) and slit diaphragm (blue)) was then used to generate a 3D model for the GBM, the central actin cables and the slit diaphragms (Fig. 5j, k and l, Mov 6). Collectively, these data show that we are able to use a machine learning model to segment the actin structures in healthy podocytes.

**Figure. 5.**
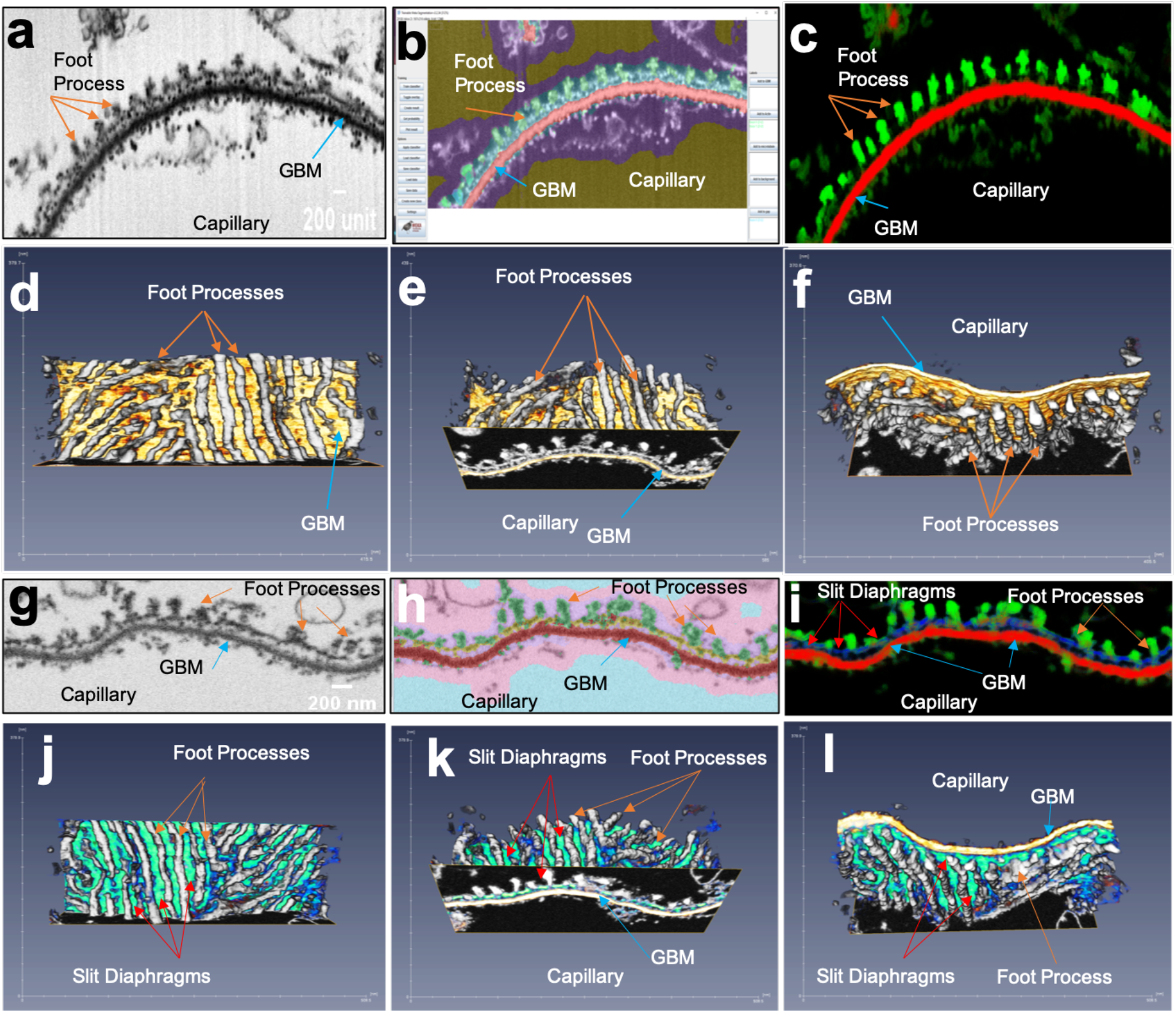
Deep learning-mediated segmentation of membrane-extracted healthy podocyte FIB-SEM images reveals the architecture of the capillary wall. (**a**) Inverted single-frame FIB-SEM image of a membrane-extracted healthy glomerulus shows the GBM and the central actin cables of the foot processes. (**b**) Example of the training process for deep learning segmentation of the image in (**a**) shows 5 classes assigned to a selected training stack (15 images). (**c**) The probability map of foot process (green) and GBM (red) channels shows the result of the deep learning classifier. (**d**) Front view of the 3D visualization of a healthy podocyte shows the interdigitating foot process. (**e** and **f**) Bottom and flipped views of the 3D visualization in (**d**). The GBM is yellow and the foot processes are silver. (**g**) Inverted single-frame FIB-SEM image of membrane-extracted healthy podocytes shows the GBM, the central actin cables of the foot processes, and the slit diaphragms. (**h**) Example of the training process for deep learning segmentation of the image in (**g**) shows 6 classes assigned to a selected training stack (15 images). (**i**) The probability map of foot process (green), GBM (red) and slit diaphragm (blue) channels shows the result of the deep learning classifier. (**j**) Front view of the 3D visualization of a healthy podocyte shows the interdigitating foot process with slit diaphragms. (**k** and **l**) Bottom and flipped views of the 3D visualization in (**j**). The GBM is yellow, the foot processes are silver, and the slit diaphragms are light blue. Scale: x and us GBM (Black arrows). Scale: tickmarks in the x and y axes are in nm.

### 3D image segmentation of Cd2ap^-/-^ glomeruli

Similar to the healthy glomeruli mentioned above, we applied the machine learning model to selected areas of the *Cd2ap^-/-^* FIB-SEM image stack (Fig. 6a and b). Unlike the WT control, training the *Cd2ap^-/-^* model required several more rounds of regional selection and the addition of other classes that separate microtubules from actin in the effaced areas. This is because the intermediate filaments and actin condensations were very close to each other, which forced us to train the software to find the clear-cut boundary segmentation between the two.

**Figure. 6.**
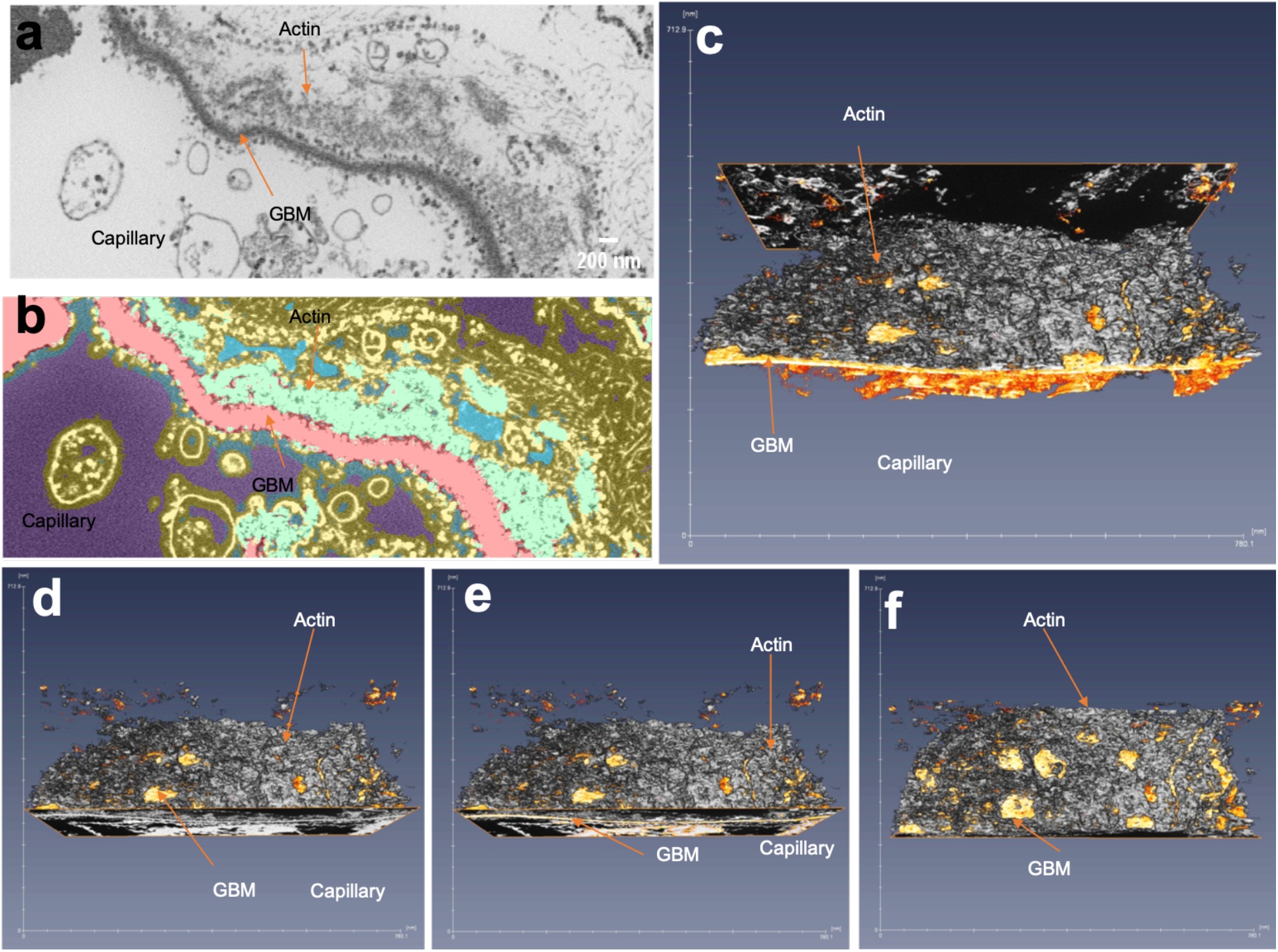
Deep learning segmented model of a membrane-extracted *Cd2ap^-/-^* podocyte shows extensive actin condensations in the effaced areas. (**a**) Inverted single-frame FIB-SEM image of a membrane-extracted *Cd2ap^-/-^* glomerulus shows the effaced area with electron-dense actin condensations near the GBM. **(b)** Example of the training process for deep learning segmentation of the image in (**a**) shows 5 classes assigned to a selected training stack (15 images). (**c**) Bottom view of the 3D visualization of a *Cd2ap^-/-^* glomerulus shows actin condensations above the GBM. (**d, e** and **f**) 3D visualizations similar to the one in (**c**) but at different angles. These views show the relative 3D position of the actin condensations with respect to the GBM. Scale: tickmarks in the x and y axes are in nm.

After multiple rounds of sequential machine learning sessions, we generated a training model for both the GBM (Fig. 6b, red) and the actin assembly in the effaced areas of the injured podocytes (Fig. 6b, light green). We then applied the trained model to a larger FIB-SEM stack (251 image stack; Mov 7 and 8), which revealed the 3D actin assembly in the *Cd2ap^-/-^* mice, showing as an extensive and dense network of actin in the effaced areas above the GBM (Fig. 6c, d, e and f, Mov 9).

### 3D image segmentation of Lamb2^-/-^ glomeruli

Consistent with previous findings [39], membrane-extraction imaging of *Lamb2^-/-^* mouse glomeruli showed segmental splitting of the GBM (Fig. 4c, 7a). Similar to the approach mentioned above for WT and *Cd2ap^-/-^* mice (Fig. 5 and 6), the machine learning approach was able to segment the split GBM and generate a trained model (Fig. 7b). Due to the splitting of the *Lamb2^-/-^* GBM, more rounds of training were needed for segmenting the split GBM as one unit while avoiding overfitting it with the actin condensations in the effaced areas. Applying this model to a large image stack (201 frames; Mov 10 and 11) allowed us to generate the first 3D model of the actin assembly inside the effaced areas of *Lamb2^-/-^* podocytes (Fig. 7c, d, e and f; Mov 12). This revealed an extensive network of actin assembly juxtaposed to the GBM.

**Figure. 7.**
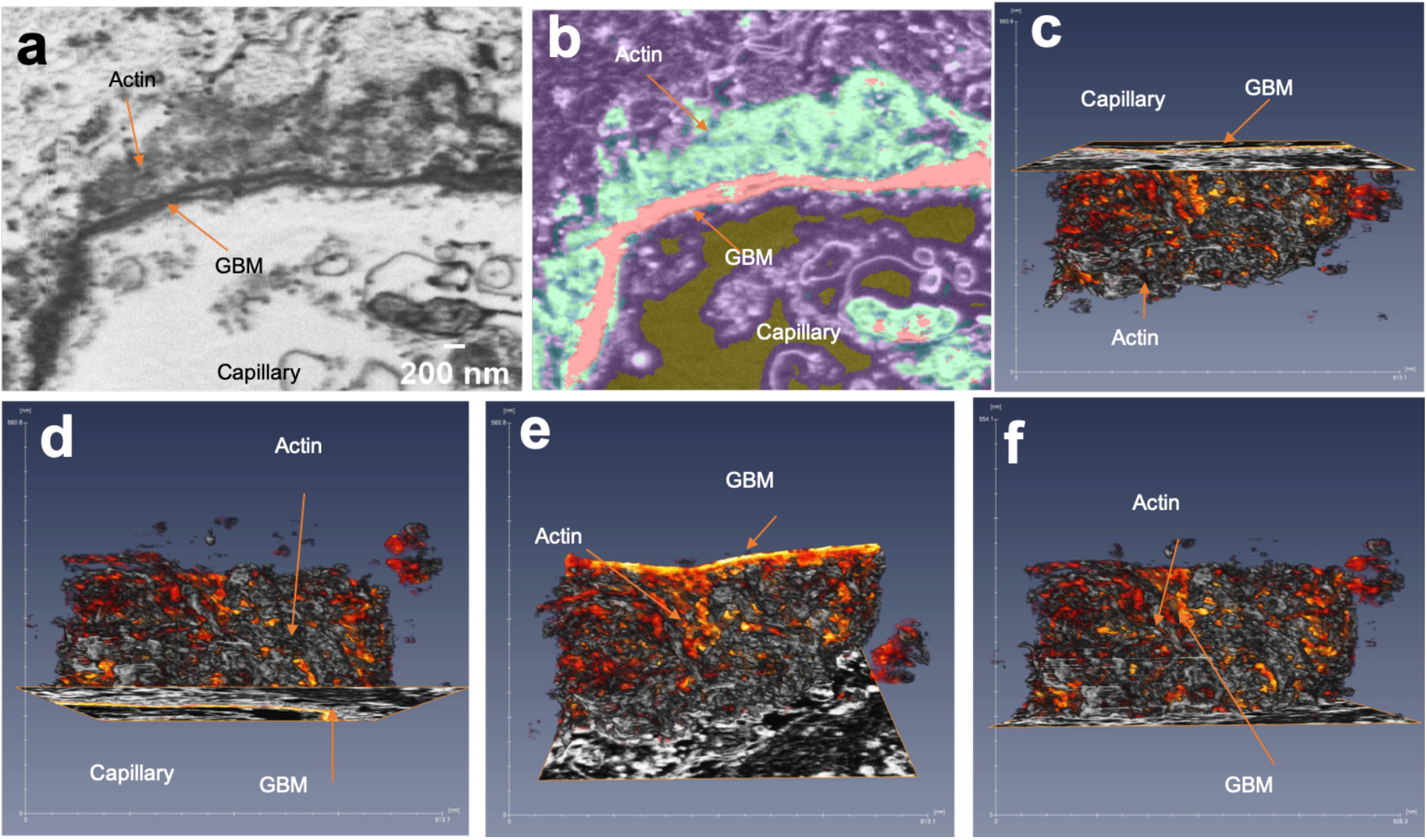
Deep learning segmented model of a membrane-extracted *Lamb2^-/-^* podocyte shows occasional GBM splitting and extensive actin condensation in the effaced areas. (**a**) Inverted single-frame FIB-SEM image of a membrane-extracted *Lamb2^-/-^* glomerulus shows the effaced area with electron-dense actin condensations near the GBM. Note the split GBM. **(b)** Example of the training process for deep learning segmentation of the image in (**a**) shows 5 classes assigned to a selected training stack (15 images). (**c**) Bottom view of the 3D visualization of a *Lamb2^-/-^* glomerulus shows the actin condensation above the GBM. (**d, e** and **f**) 3D visualizations similar to the one in (**c**), but at different angles. These views show the relative 3D position of the actin condensation in reference to the GBM. Scale: tickmarks in the x and y axes are in nm.

### 3D image segmentation of Alport glomeruli

As observed before [23], membrane-extracted Alport glomeruli (60 days of age) showed grossly irregular GBMs with extensively effaced foot processes (Fig. 4d and Fig. 8a; Mov 11 and 12). The GBM showed much heterogeneity between the podocyte and endothelial cell leaflets, but not the typical basket weave appearance observed by TEM (Fig. 8a, f and g, blue arrows [55]). Similar to the other disease models in this study (i.e., *Cd2ap^-/-^* and *Lamb2^-/-^*), podocytes were effaced and showed widespread electron dense material juxtaposed to the GBM (Fig. 8a, f and g, orange arrows). Because of the complexity of this sample, we manually segmented the GBM, which revealed an irregular 3D structure (Fig. 8d and e). Machine learning for segmentation of the actin cytoskeleton in the effaced areas was challenging because of the similar intensities of the irregularly thickened GBM and the electron dense patterns inside the effaced areas. We therefore manually broke down the image stacks (150 frames, Fig. 8g, Mov 13 and 14) into the GBM (Fig. 8h) and effaced podocyte areas (Fig. 8i). Training the GBM classifier required recognition of not only the inner and outer leaflets of the GBM, but also the region in between. To train the algorithm to recognize podocyte actin condensations, we selected regions similar to those in the *Cd2ap^-/-^* stack, then built a complete 3D reconstruction of the structures of interest (j, k and l).

**Figure. 8.**
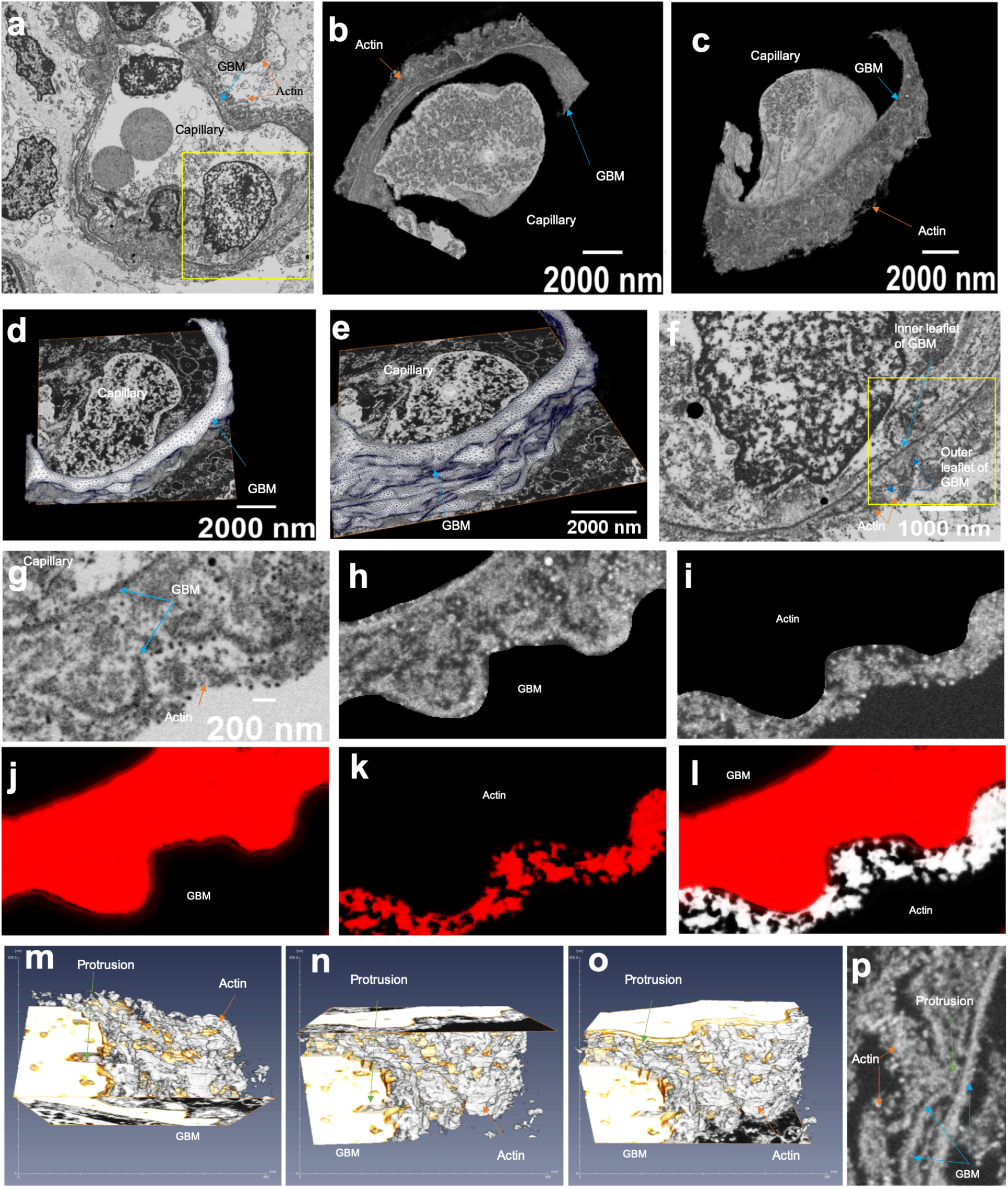
Manual and deep learning segmentation of a *Col4a3^-/-^* Alport membrane-extracted glomerulus shows irregularly thickened GBM and extensive actin condensations in podocytes. (**a**) Inverted single-frame FIB-SEM image of a membrane-extracted Alport glomerulus shows the irregularly thickened GBM and podocyte FPE with actin condensations near the GBM. Blue arrows indicate the GBM while orange arrows indicate the actin condensations. (**b** and **c)** 3D rendering of the ROI on the image stack in (**a**) shows the actin condensations near the thickened GBM. (**d** and **e**) 3D renderings of manually segmented GBM of the same ROI in (**a**) show the irregular bumpy nature of the Alport GBM. (**f**) Zoomed-in image of (a) shows the actin condensation above the irregularly thickened GBM. The boxed area is the ROI of the FIB-SEM image stack used for deep learning. Note the heterogeneous material between the two GBM leaflets (blue arrows). (**g**) Single frame of the FIB-SEM image stack of the area in (**f**) before machine learning segmentation. (**h** and **i**) Manual breakdown of the area in (**g**) into GBM (**h**) and actin condensation (**i**) before using them for deep learning. (**j**) Probability map of the GBM after applying the classifier on the full stack in (**h;** 150 frames) shows the thickness between the two leaflets of the GBM. (**k**) Probability map of the actin condensation after applying the classifier on the full stack in (**i;** 150 frames) shows that deep learning segmentation is able to identify the detailed actin condensation. (**l**) Overlay of the two probability maps in (**j**) and (**k**) shows the final two channels map that is used for 3D reconstruction. (**m, n** and **o**) Three angles of the 3D reconstruction of an Alport glomerulus show the actin assembly in the effaced area atop the irregularly thickened GBM. Note the actin protrusions into the GBM (green arrows). (**p**) Corresponding single FIB-SEM frame image of the image stack processed in (**m**, **n** and **o**) shows the electron-dense actin protrusion embedded in the GBM (green arrow). Scale: tickmarks in the x and y axes are in nm.

The resulting trained model for identifying the GBM and actin condensations was applied to the larger image stack to generate a 3D model of the Alport GBM and of the thickened, irregular actin cables in the effaced areas (Fig. 8m, n and o, Mov 15). In some areas, actin protrusions embedded in the GBM were evident (Fig. 8m, blue arrows in n, o and p; Mov 15), consistent with the earlier report showing podocyte process invasion into the Alport GBM [23].

### Intermediate filaments in the membrane-extracted glomeruli

To gain more insights into the 3D organization of the intermediate filaments that were observed in the WT controls (Fig. 1i, j, k; Fig. S2 a, b and c), we applied a deep learning approach to segment those structures (Fig. S2d) and render them in 2.5D (Mov 16) and 3D (Fig. S2e, f and g; Mov 17). Results were consistent with earlier immuno-gold staining observations that major processes contain loosely organized intermediate filaments [56, 57]. In contrast, injured podocytes presented much denser and compact sets of intermediate filaments in the effaced areas above the actin assembly in all our studied injury models (Cd2ap-/-: Fig. S2 h, I, j and k; Mov 18 and 19; Lamb2 -/-: Fig. S2 l, m, n and o; Mov 20 and 21; Alport syndrome: S2 p, q, r and s; Mov 22 and 23).

## Discussion

Our technique provides a new method to visualize the actin and intermediate filament networks of podocytes in their native environment, the glomerulus. This approach has provided interesting new observations about the podocyte’s ultrastructure:

i. The central actin cables within foot processes are connected to, and appear continuous with, the slit diaphragms. This is consistent with previous observations that predicted this connection biochemically [58, 59]. It is interesting that the mild detergents used here (i.e., 0.01% saponin and 0.5% Triton-X100) did not disrupt the slit diaphragm assembly, in agreement with the finding that nephrin and podocin are solubilized only when the actin cytoskeleton is disassembled [58].
ii. Together, the actin cables within foot processes and the slit diaphragms to which they connect form one continuous, mesh-like, electron dense sheet that covers the GBM in healthy cells. This network should act as a unit that is capable of resisting the pulsating forces generated across the capillary wall from both the intracapillary blood pressure against the wall and the shear stress of fluid flow through the filtration slits. Our observations are consistent with the GBM compression model proposed by Miner and Fissell [60] in which foot processes linked with one another via slit diaphragms act together to counterbalance the hydrodynamic forces across the glomerular capillary wall; the resulting compression of the GBM leads to pores sizes that restrict molecules larger than 65 kDa (such as albumin) from passing through to reach the urinary space.
iii. Podocyte cell bodies and the major processes are reinforced by an assembly of thick parallel actin bundles that run along their longitudinal axes. Based on 1) the similarities between the actin cables imaged by FIB-SEM (i.e., viewed in 2D as actin patches, Fig. 3) and the synaptopodin-positive patches at the periphery of the podocyte cell body and the major processes observed by STORM [31]; and 2) the localization of myosin IIA high in the podocytes (i.e., in the cell body and major processes but not in foot processes [31]), we speculate that these actin bundles observed by FIB-SEM are contractile in nature, which generate tension necessary for maturation of podocyte adhesive complexes at the GBM, analogous to tension thresholds for adhesion of other cells [61–63].
iv. After injury, podocytes undergo dramatic shape changes as foot processes become effaced, and this coincides with extensive actin rearrangements and the appearance of ectopic, irregularly thickened basal actin assemblies in the effaced areas. This is consistent with models of foot process fragility arising from cytoskeletal instability [64, 65]. The three mouse models of podocyte injury and nephrotic range proteinuria that we studied have different manifestations and mechanisms of injury. The loss of CD2AP affects the stability of slit diaphragms and their linkage to the actin network [38], whereas *Lamb2^-/-^* and *Col4a3^-/-^* Alport mice affect the laminin and collagen IV trimer networks in the GBM, respectively [39, 40]. The striking similarities in actin network architecture that we observed basally in the effaced areas in these models suggests that common pathways govern the cytoskeletal changes leading to FPE.

Since its description 25 years ago [30], the prevalence and significance of the injured podocyte’s “actin mat” has not been well documented. Here, we show an extensive actin assembly that is reminiscent of the actin mat; in 3D it presents as randomly organized, thickened actin cables with random directions in-plane. We found this actin assembly in three animal models with severe FPE. Whether these changes in the actin network are always present in FPE is still an open question, and we believe that our membrane-extraction technique can answer it. For this study, we chose the mouse models at an age when foot processes are massively effaced, but it would be interesting to assess the same mice at earlier ages or similar mice but with milder kidney phenotypes (such as *Lamb2*^-/del44^ mice [66]).

Although we tried to minimize artifacts due to the detergent-extraction process by early fixation and extensive washes, we did observe some vacuolization in the major processes, cell body, and the surrounding areas, although they did not disturb the cytoskeletal network. To minimize disruption to the cytoskeleton’s architecture, we stabilized the microfilaments using phalloidin and taxol. Despite the 1μM taxol treatment, extensive microtubule networks were not observed, possibly suggesting that any attachment between the microtubules and the remaining cytoskeletal elements was relatively weak following fixation. However, the extensive network of intermediate filaments suggests that these intermediate filaments were stable and well connected to the actin cytoskeleton, as recently suggested [67]. The intermediate filaments might possibly be vimentin [56, 68], nestin [69–71] or desmin [72], as early reports suggested an upregulation in these proteins in injured podocytes [57, 72, 73]. We hope to identify these through future FIB-SEM studies with membrane-extracted glomeruli from mice lacking these proteins.

This study is the first of its kind that successfully uses a deep learning approach in combination with FIB-SEM imaging. Since manual segmentation of podocytes requires time and introduces bias [26, 27], a machine learning approach was critical. Amongst the many approaches available to facilitate segmentation of biological samples by machine learning [74], we chose the open-source Weka in Fiji/ImageJ [52] because of its simplicity, speed, and flexible use of filters for machine learning. When performing our deep learning segmentation, we had to choose whether to create one single classifier that roughly segments all classes of all datasets, or create separate classifiers that segment each dataset independently [74]. Given the complexity of our datasets, the separate time for sample preparation and imaging for each of them, and internal differences observed across the four biological samples that we used, applying one single deep learning classifier model to all injury models resulted in loss of accuracy. Although several strategies exist to compensate for this problem of “overfitting” [18], we achieved satisfactory segmentation by generating classifiers for each of our datasets. The approach allowed for the supervised segmentation of podocyte structures with high accuracy as compared to manual segmentation, the latter validated by choosing smaller datasets from each condition and validating results using manual segmentation. The result was a fast, reliable 3D reconstruction of large FIB-SEM image stacks. This approach, when integrated with membrane extraction, is a general framework that we believe will have applicability to cells in diverse organs in cases where actin cytoskeletal structures are dependent upon the cells’ mechanical microenvironment.

## Supporting information

Overview WT images stack

WT image stack of the major processes

3D visualization of the actin cables in the major processes

WT image stack of the actin in the foot processes

3D segmentation of WT showing actin cables in foot processes

3D segmentation of WT showing slit diaphragms between the foot processes

Overview Cd2ap KO images stack

Cd2ap KO image stack region of interest showing the actin condensation in the effaced area

3D segmentation of Cd2ap KO showing the actin condensation in 3D

Overview Lamb2 KO images stack

Lamb2 KO image stack region of interest showing the actin condensation in the effaced area

3D segmentation of Lamb2 KO showing the actin condensation in 3D

Overview Col4a3 KO images stack

Col4a3 KO image stack region of interest showing the actin condensation in the effaced area

3D segmentation of Col4a3 KO showing the actin condensation in 3D

2.5D rendering of WT stack showing the intermediate filaments

3D segmentation of WT stack showing the intermediate filaments

2.5D rendering of Cd2ap KO stack showing the intermediate filaments

3D segmentation of Cd2ap KO stack showing the intermediate filaments

2.5D rendering of Lamb2 KO stack showing the intermediate filaments

3D segmentation of Lamb2 KO stack showing the intermediate filaments

2.5D rendering of Col4a3 KO stack showing the intermediate filaments

3D segmentation of Col4a3 KO stack showing the intermediate filaments

## Acknowledgments

We acknowledge the assistance of Matt Joens, Dennis Oakley, Dr. Sanja Sviben, Dr. Praveen Krishnamoorthy, and Dr. James Fitzpatrick for microscopy and imaging analysis performed at the Washington University Center for Cellular Imaging (WUCCI), which is supported by Washington University School of Medicine, The Children’s Discovery Institute of Washington University and St. Louis Children’s Hospital (CDI-CORE-2015-505 and CDI-CORE-2019-813), the Foundation for Barnes-Jewish Hospital (3770 and 4642), and the Washington University Diabetes Research Center (NIH P30 DK020579). This work was also supported by grants from the American Heart Association (17SDG33420069 to HYS), the NIH (P30DK020579 to HYS and R01DK058366 and R01DK078314 to JHM), the NSF Science and Technology Center for Engineering MechanoBiology (CMMI 1548571), the Office of the Vice Chancellor for Research at Washington University, and the Division of Nephrology at Washington University School of Medicine.

---

**Figure. S1.**
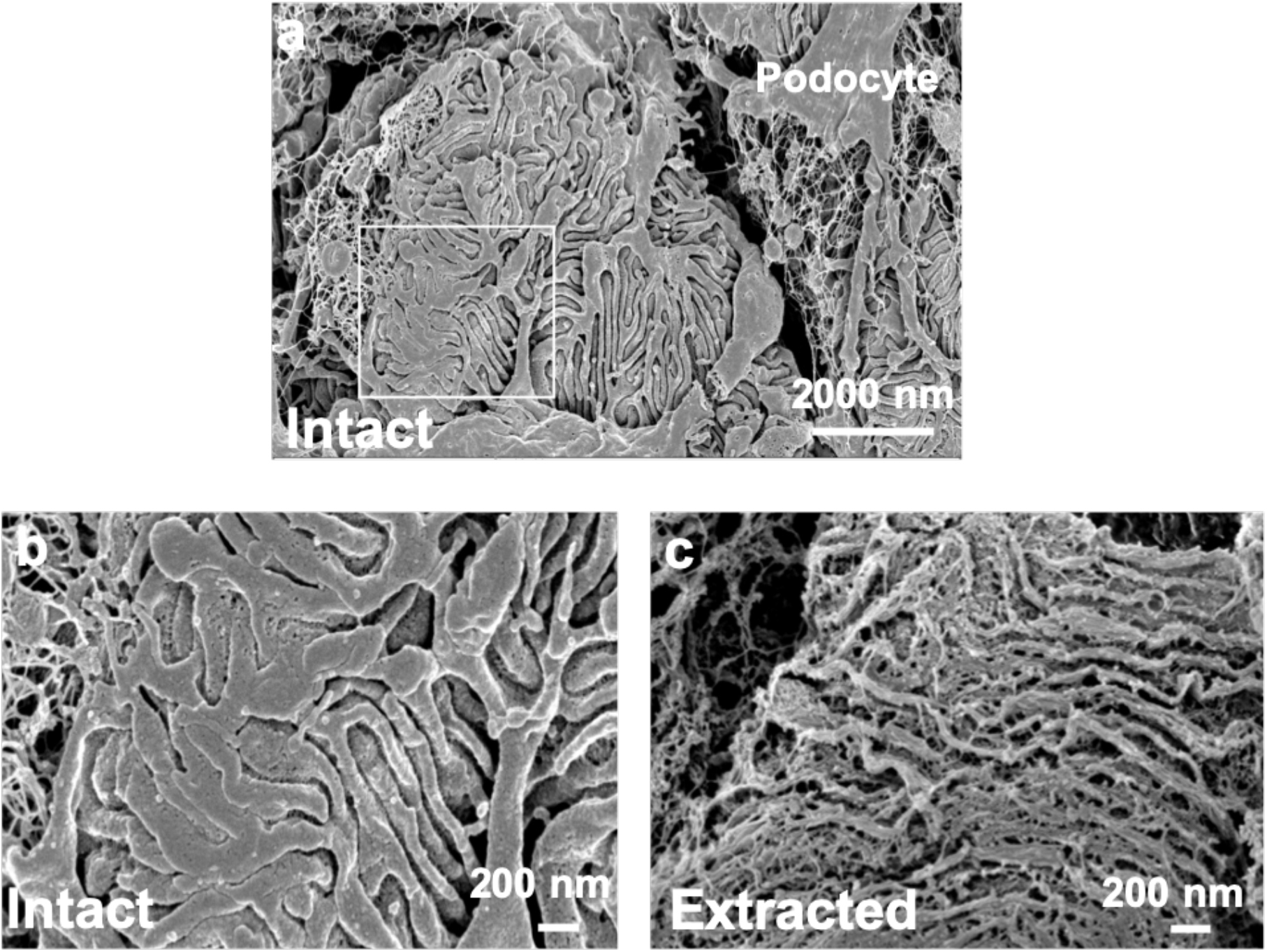
SEM images of intact and membrane-extracted *Lamb2^1^* glomeruli show the widening of foot processes as a sign of foot process effacement (FPE) before and after cell membrane removal. (**a** and **b**) Low and high magnification images, respectively, of an intact *Lamb2^-/-^* glomerulus show FPE. (**c**) A membrane-extracted *Lamb2^-/-^* glomerulus shows irregularly distributed actin bundles covering the capillary wall.

**Figure. S2.**
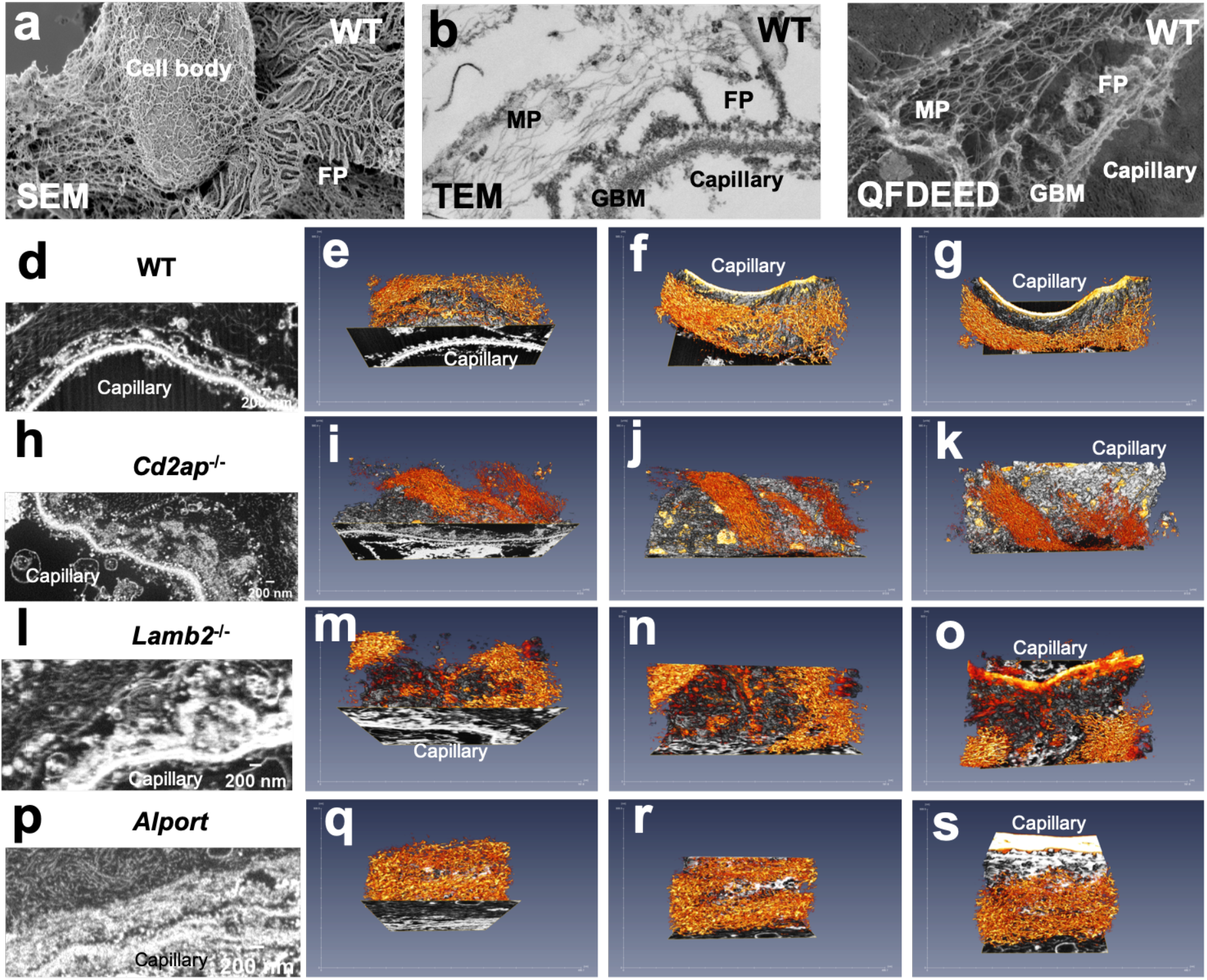
Fine podocyte microfilaments in membrane-extracted glomeruli appear irregular using deep learning segmentation. (**a**) SEM image of a membrane-extracted WT glomerulus shows microfilaments extending from the podocyte nuclear region towards the foot process (FP) areas. (**b** and **c)** Transmission (TEM) and quick-freeze deep-etch EM (QF-DEEM) micrographs show WT podocytes with microfilaments located in the major processes (MP) but not the foot processes (FP). (**d**, **h**, **l** and **p**) Single slices of the FIB-SEM image stacks (inverted) of all the conditions used for deep learning for the microfilament segmentation. (**e**, **f** and **g**) Three angles of the 3D reconstruction of the WT glomerulus in (**d**; 251 frames) show the microfilament assembly (orange) relative to the actin cables and slit diaphragms in the foot processes. (**i**, **j** and **k**) Three angles of the 3D reconstruction of the *Cd2ap^-/-^* glomerulus in (**h**, 251 frames) show the microfilament assembly (orange) relative to the actin assembly in the effaced foot processes. (**m**, **n** and **o**) Three angles of the 3D reconstruction of the *Lamb2^-/-^* glomerulus in (**l**, 251 frames) show the microfilament assembly (orange) relative to the actin assembly in the effaced foot processes. (**q**, **r** and **s**) Three angles of the 3D reconstruction of the podocyte area in the Alport glomerulus in (**p**, 251 frames) show the microfilament assembly (orange) relative to the actin assembly in the effaced foot processes. Scale: tickmarks in the x and y axes are in nm.

